# Plant Molecules Protect Against Inflammatory Bowel Disease by Restoring Gut Microbiota-Immune Homeostasis and Suppressing Pro-inflammatory Markers

**DOI:** 10.64898/2026.08.24.745385

**Authors:** Karma Yeshi, Subir Sarker, Md. Zohorul Islam, Darren Crayn, Stephen G. Pyne, Paul Giacomin, Matt Field, Md. Mizanur Rahaman, David Wilson, Michael J. Smout, Norelle L. Daly, Alex Loukas, Roland Ruscher, Phurpa Wangchuk

## Abstract

Inflammatory bowel disease (IBD) is associated with chronic intestinal inflammation and gut microbial dysbiosis, yet effective microbiome-targeted therapeutics remain limited. Here, we investigated the anti-inflammatory and microbiome-modulating activities of metabolites isolated from *Garcinia brassii*, an endemic species of the Australian Wet Tropics. Five compounds, including a new natural product named garcitine, were isolated and structurally characterised. In human immune cells, garcinol and garcinia biflavonoid 1 significantly suppressed lipopolysaccharide-induced production of IL-1β, IL-6, and TNF without detectable cytotoxicity, while parvifoliol F selectively inhibited IL-1β release. Therapeutic efficacy was further evaluated in a TNBS-induced murine colitis model, where garcinia biflavonoid 1 and parvifoliol F significantly reduced colonic inflammation and improved histopathological outcomes. 16S rRNA sequencing demonstrated that both compounds restored gut microbial homeostasis by reversing colitis-associated dysbiosis and reducing inflammation-associated microbial signatures. Functional pathway prediction further suggested suppression of pro-inflammatory microbial metabolic pathways following treatment. Together, these findings demonstrate that *Garcinia*-derived metabolites alleviate experimental colitis through coordinated immunomodulatory and microbiome-reprogramming mechanisms and identify garcinia biflavonoid 1 and parvifoliol F as promising candidates for microbiome-targeted IBD therapeutics.

## 1. Introduction

Inflammatory bowel disease (IBD) is a lifelong chronic disease that affects more than 5 million people globally(Burisch et al., 2023), with increasing incidence in newly industrialized countries(Prideaux et al., 2012). IBD has two subtypes, namely ulcerative colitis (UC, affecting the colon) and Crohn’s disease (CD, affecting any part of the gastrointestinal tract)(Rahaman et al., 2025; Zhang and Li, 2014). UC is driven by the response of T helper 2 (Th2) cells, characterized by IL-5 and IL-13 produced by natural killer T (NKT) cells. CD, in contrast, is more Th1/Th17 cell-driven, with IL-12 and IL-23 as major cytokines(Strober and Fuss, 2011). The exact etiology of IBD is unknown and is said to be influenced by multiple factors, including genetic susceptibility, immune factors, and the gut microbiota(Rahaman et al., 2025; Schaubeck et al., 2016; Wangchuk et al., 2024). Increasing evidence suggests a major role for gut microbiota dysbiosis in driving IBD(Kaur et al., 2011; Oligschlaeger et al., 2019). Among beneficial gut microbiota, those capable of producing short-chain fatty acids (SCFAs), such as *Faecalibacterium prausnitzii* help protect against IBD by producing short-chain fatty acid (SCFA) – butyrate(Visconti et al., 2019), gut barrier integrity(Ghosh et al., 2021), and motility(Waclawiková et al., 2022). Butyrate is associated with protection against IBD by inhibiting activation of NF-κB, enhancing regulatory T cell function, and promoting expression of tight junction proteins(Ambat et al., 2024). Prebiotics enriched with butyrate-producing bacteria have been shown to promote a healthy gut microbiome(Breuer et al., 1997; Scheppach et al., 1992). Currently, there is no cure, and existing treatments can only manage IBD(Faubion et al., 2001). Moreover, existing drugs have adverse side effects and often become ineffective over time(McLean and Cross, 2014). Due to the enormous disease burden, there is an urgent need for new and effective treatments with fewer side effects.

Natural resources, including medicinal plants, have long been a major source of drugs, and they have proven effective against many diseases, including IBD(Newman and Cragg, 2020). For instance, *Garcinia* (family Clusiaceae) species are widely known for their anti-inflammatory compounds(Hemshekhar et al., 2011), such as polyisoprenylated polyphenols(Krishnamurthy et al., 1981). Infusions prepared from *Garcinia* fruits are popular for treating IBD-related conditions, including ulcers and dysentery, in traditional medicine systems of many countries(Acuña et al., 2012). Moreover, crude extracts and pure compounds from *Garcinia* species have significantly improved gut microbiome homeostasis(Gutierrez-Orozco et al., 2015; Kang et al., 2023). For instance, treatment with α-mangostin, isolated from other *Garcinia* species have been found to reduce the abundance of harmful *Firmicutes* and increase the abundance of beneficial *Proteobacteria* in mice(Gutierrez-Orozco et al., 2015; Yang et al., 2024).

In this study, we have investigated the anti-inflammatory properties of three compounds derived from *Garcinia brassii* (endemic to the Wet Tropics of Australia) in an acute colitis mouse model showed that some of these compounds restored colitis-induced dysbiosis. Our findings support the potential of these compounds as new therapeutic agents for managing IBD through enhancing gut microbiome homeostasis.

## 2. Materials and methods

### 2.1 Isolation and characterisation of compounds

The 80% EtOH extract of the aerial parts of *G. brassii* was purified using repeated chromatographic techniques, such as high-performance liquid chromatography (HPLC), column chromatography, and nuclear magnetic resonance (NMR) spectroscopy. Five compounds were characterised and identified using spectroscopic data. A detailed description of the isolation and characterisation of five compounds is given in the supplementary information.

### 2.2 Compound preparation for cell toxicity assessment and immunoassays

Compounds **1**–**4** (1 mg) were dissolved in 10 µL of dimethyl sulfoxide (Sigma-Aldrich, St. Louis, MO, USA) and subsequently diluted and adjusted to a final concentration of 1 mg/mL with 990 µL of RPMI-1640 media (without L-glutamine), Gibco Thermo Fisher Scientific, Waltham, MA, USA. The desired concentrations were prepared from this stock solution for cell viability testing.

### 2.3 Human PBMC collection and culture conditions

Human PBMC were isolated from buffy coats of healthy donors (n=1-3) using a lymphoprep™ density gradient medium (STEMCELL Technologies, Vancouver, Canada) as per the product protocols under ethics (ethics no. H8523) approved by the James Cook University human research ethics committee. The PBMC assay culture was conducted as per our previous studies(Ritmejeryte et al., 2022; Yeshi et al., 2022).

### 2.4 Compound screening for cell viability using the xCELLigence assay

Before testing for anti-inflammatory activity, the purified compounds **1**–**4** were tested for their effects on the human bile duct epithelial (h69) cells monitored with the xCELLigence assay. The xCELLigence, RTCA, Single Plate System (ACEA Biosciences, Inc., Santa Clara, California) measures cell viability of the human bile duct epithelial (h69) cells treated with 10 µg/mL of the compound relative to a DSMO control in real-time. A real-time and label-free xCELLigence system monitored cells’ health for 72 h. The xCELLigence system has microtiter plates attached with digitalised gold microelectrodes, which continuously monitor the viability of cultured cells through electrical impedance/perturbations as the readout, making it possible to distinguish between senescence, cell toxicity (cell death), and reduced proliferation (cell cycle arrest). The data shown are pooled data plotted as averages with SEM bars from 2 experiments of 5 replicates. Averaged data were compared against DMSO control at each time point, with 2-way ANOVA and Holm-Sidak’s multiple comparisons.

### 2.5 Proinflammatory cytokines suppression assay and quantification

Briefly, 1 × 10^6 cells in 100 μL of R-10 medium were seeded into each well of a 96-well U-bottom plate (Falcon, Corning, USA). The R-10 medium consisted of RPMI-1640 (Gibco) supplemented with 10% heat-inactivated fetal bovine serum (FBS; Corning), penicillin (100 U/mL), and streptomycin (100 μg/mL) (Gibco). PBMCs were stimulated with either lipopolysaccharide (LPS, 10 ng/mL; Sigma-Aldrich) or a combination of phorbol 12-myristate 13-acetate (50 ng/mL) and ionomycin (1 μg/mL) (P/I; eBioscience), followed by incubation at 37 °C in 5% CO2 for 2 h. Subsequently, pure compounds, dexamethasone, or R10-DMSO (10 μg/mL) were added to the corresponding wells in triplicate. The cells were then incubated overnight at 37 °C in a humidified atmosphere containing 5% CO2. The following day, plates were centrifuged at 277 × g for 5 min at 4 °C, and culture supernatants were harvested and stored at −80 °C until cytokine assessment. Cytokine concentrations were measured using the LEGENDplex™ Human Inflammation Panel 1 (Catalogue no. 740808; lot no. B291817, BioLegend®, USA), which enables simultaneous detection of 13 inflammatory cytokines, including IL-1β, IFN-α2, IFN-γ, TNF, MCP-1, IL-6, IL-8, IL-10, IL-12p70, IL-17A, IL-18, IL-23, and IL-33. Samples were analysed using an LSRFortessa flow cytometer (BD), and data were processed with LEGENDplex™ software (version 2022-02-10, Qognit, Inc.). Results are presented as mean ± standard error of the mean (SEM).

### 2.6 Mice

BALB/c mice (age-matched, 5-wk-old male) were purchased from the ARC (animal resource centre) in Perth, Australia. Mice were randomly distributed on arrival into groups of 5 per cage in all experiments to maintain consistency. In two consecutive experiments, 10 mice in each treatment group were used to ensure statistical significance. Mice were allowed to acclimatize for at least five days in the animal facility room before any experiment. Throughout the experiments, mice were kept at an optimal temperature (20-26 °C), humidity (40-70%), and a 12-hour day/night cycle. They were fed sterile mouse chow purchased from Specialty Feeds of Glen Forrest (Western Australia) and reverse osmosis (RO) water ad libitum. The cage was changed weekly to minimize the stress. All animal studies were carried out under the ethics (Ethics no. A2647) approved by the animal ethics committee of James Cook University.

### 2.7 TNBS-induced colitis and treatment with compounds

Experimental colitis was established in mice using a TNBS-induced model as previously reported by Wangchuk et al.(Wangchuk et al., 2015). Animals were anaesthetised by intraperitoneal (IP) injection of ketamine (6.6%) and xylazine (0.6%) diluted in phosphate-buffered saline (PBS) (Provet). Following anaesthesia, colitis was induced on day 0 through intrarectal (IR) administration of 2.5 mg TNBS (Sigma) prepared in 100 µl of 50% ethanol. On day 1, mice assigned to treatment groups received either the test compound or dexamethasone (20 µg in 200 µl PBS) via IP injection. Naïve and TNBS control groups were administered the corresponding vehicle solution (200 µl DMSO/PBS) by the same route. Animals were monitored daily for body condition and clinical manifestations of colitis using previously established scoring criteria (Wangchuk et al., 2015). On day 4, mice were euthanised by CO₂ inhalation, and colons were collected for macroscopic assessment of inflammation and measurement of colon length.

### 2.8 Genomic DNA extraction and full-length 16S sequencing

Genomic DNA was extracted from archived mouse colon samples using the DNeasy PowerSoil Pro Kit (Qiagen, Germany) according to the manufacturer’s instructions with slight modifications. Briefly, intestinal mucosal material was aseptically scraped using a sterile scalpel and suspended in PBS in sterile microcentrifuge tubes. Mechanical homogenisation was performed by bead beating with a TissueLyser-LT2 instrument (Qiagen, Germany) for approximately 10 min to ensure effective tissue disruption. Samples were subsequently centrifuged at 8,000 × g for 5 min, and the clarified supernatant was processed for DNA isolation following the standard kit protocol. DNA concentration and purity were determined using a NanoDrop spectrophotometer, and samples with concentrations above 50 ng/µl and A260/A280 ratios between 1.8 and 2.0 were selected for downstream sequencing analyses. Amplification of the full-length bacterial 16S rRNA gene and primer optimisation were performed by the Australian Genome Research Facility (AGRF, Brisbane, Australia). Full-length 16S sequencing was employed to achieve improved taxonomic resolution and more accurate profiling of microbial community composition. Indexed amplicons were pooled and sequenced using PacBio SMRTbell long-read sequencing technology at AGRF.

### 2.9 Microbiome sequence data analysis

PacBio HiFi full-length 16S rRNA sequence data were processed using the QIIME2 workflow, with denoising and quality filtering conducted through the DADA2 algorithm to generate high-confidence amplicon sequence variants (ASVs). Taxonomic classification was performed using two complementary strategies. First, ASVs were assigned using VSEARCH consensus alignment against the Genome Taxonomy Database (GTDB release 207) to maximise taxonomic consistency. In parallel, a naïve Bayesian classification approach implemented in DADA2 was applied using a hierarchical reference database system. This sequential workflow queried GTDB r207, SILVA v138, and the NCBI RefSeq 16S rRNA database supplemented with the Ribosomal Database Project (RDP), thereby enhancing identification of low-abundance or poorly represented taxa. Microbial α-diversity was evaluated using Observed richness, Shannon diversity, and Simpson diversity indices. β-diversity analyses were conducted using unweighted UniFrac distance matrices. Ordination analysis based on Bray–Curtis dissimilarity was performed using principal component analysis (PCA) within the Phyloseq package (v1.48) (McMurdie and Holmes, 2013) in R. Differential abundance analysis was undertaken using DESeq2 (v1.44), whereas data visualisation was generated using ggplot2 (v3.5.1) (Wickham, 2016) and pheatmap v1.0.12(Kolde, 2019). Statistical analyses were conducted using base R software. Functional inference of microbial metabolic pathways was predicted using PICRUSt2 (Douglas et al., 2020) in combination with the MetaCyc database.

### 2.10 Human LEGENDScreen assay

Cryopreserved PBMC isolated from buffy coats of three healthy donors (BC970050, BC803935, and BC814304) were thawed, washed, and counted as described above. After a washing step, cells from three different donors were seeded into a 6-well flat-bottom plate (∼12.5 × 10^6^ cells per well) and stimulated with a cocktail of 50 ng/mL PMA and 1μg/mL ionomycin (P/I; eBioscience) and incubated at 37 ^◦^C, 5.5% CO_2_ for 2 h. After 2 h post-stimulation, pure compounds (**2** and **3,** 10 μg/mL)/R10-DMSO were added into the respective wells. The plates containing all treatments were incubated overnight in a CO_2_ incubator (5% CO_2_ and 37 °C).

Cells from each condition from each donor were pooled and stained with the CD45 antibodies in four different fluorochromes (PERCP-CY5.5, BUV395, FITC, and APC) for 30 min on ice in the dark. Following staining, cells were washed three times with staining buffer, and equal numbers of cells from individual donors were pooled. Subsequently, 3 × 10^5 cells were dispensed into each well of the LEGENDScreen Human Cell Screening Kit plates (BioLegend, 700001), which contained 25 μL of antibody solution with a unique antibody specificity in each well. The plates were prepared in accordance with the manufacturer’s instructions and maintained at 4 °C. After incubation on ice for 30 min, the plates were washed with staining buffer, and cell pellets were resuspended in a staining cocktail containing CD3-APC-H7, CD11c-BV510, CD19-UV395, CD69-V650, HLA-DR-PE-Cy7, CD161-BV785, CD185-PE CF594, CD25-PE, CD4-PerCP5.5, and CD8-FITC antibodies. Cells were incubated with the staining cocktail for 30 min on ice in the dark. Following washing, viability staining was performed for 15 min at room temperature using LIVE/DEAD™ Fixable Near-IR Dead Cell Stain (Invitrogen, Thermo Fisher Scientific; Lot 2652701). Cells were then washed again, fixed in 1% paraformaldehyde (PFA) for 30 min, and analysed using a BD LSRII flow cytometer. Data were analysed with FlowJo software version 9.

### 2.11 Pathway enrichment gene ontology and analyses

Gene ontology (GO) term and pathway enrichment analyses were generated using a combination of Enrichr(Kuleshov et al., 2016), DAVID(Sherman et al., 2022), R v4.2.3, and consensus software code previously described(Waardenberg and Field, 2019). Background gene lists for 342 genes were first input to DAVID and EnrichR using all *xCELLigence* panel targets found to correspond to annotated ENSEMBL genes. *xCELLigence* scores at least 1.5 times higher or lower relative to baseline levels were categorised as being up- or down-regulated, respectively. For both DAVID and Enrichr, GO terms and pathway enrichment outputs were generated, and these data were input to R, with all terms with a *p*-value of ≦ 0.05 retained. Plots were generated with the R package ggplot for enriched GO terms and Pathways with sorted *p*-values on the y-axis and further annotated if the FDR < 0.2.

### 2.12 Data Analysis

The NMR data were analyzed with MestReNova (version 14.3.1-31739, Mestrelab Research S.L., Santiago de Compostela, A Coruña, Spain). The compound structures and their 2D NMR correlations were drawn with the help of ChemDraw software (version 21.0.0). The statistical significance for cytotoxicity and cytokine release data between untreated and stimulated groups was determined by a one-sample *t*-test and Mann-Whitney U test in GraphPad Prism software (version 9.0).

## 3. Results

Since this research focuses on how natural compounds alleviate ulcerative colitis (IBD) through the restoration of the gut microbiome, the isolation and characterisation of compounds from *Garcinia brassii* have been mostly provided in the Appendices. We isolated and characterised five compounds, including one new natural product, which we named it as garcitine (**1**) and four known compounds, namely garcinol (**2**)(Pasaribu et al., 2021), garcinia biflavonoid 1 (**3**)(Han et al., 2005), parvifoliol F (**4**)(Rukachaisirikul et al., 2006), and friedelin (**5**) (Ambarwati et al., 2019; Gottlieb et al., 1985) (Chemical structure provided in SI Appendix, Fig. 1). The characterisation and spectroscopic data of compounds **1**–**5** are provided in the SI Appendix, Tables S1-S2 and Fig. S4– S14.

**Fig. 1.**
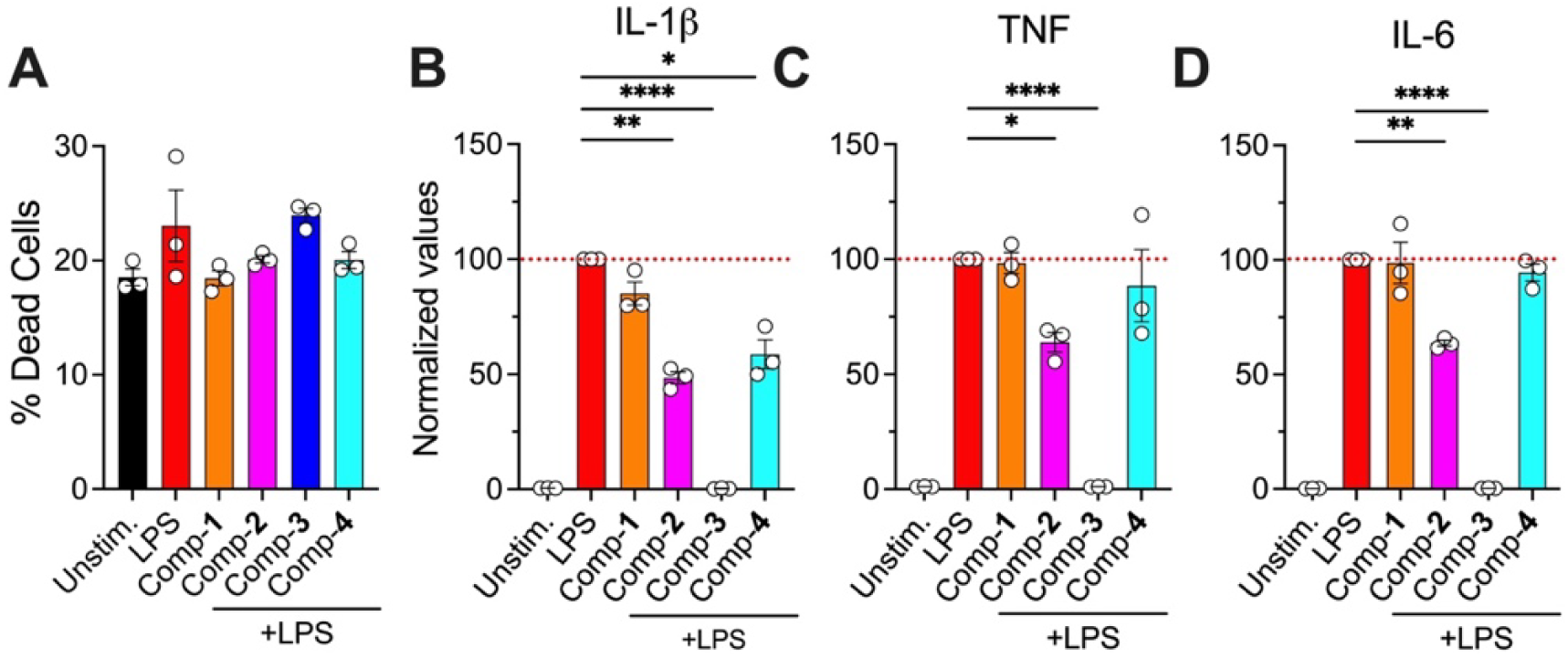
Anti-inflammatory effect of compounds (1–4). (**A**) Percentage of viability of human PBMC after overnight incubation with compounds (**1**–**4**) at 10 μg/mL, (**B**) The inhibitory effect of compounds (10 μg/mL) on the release of proinflammatory cytokines in the supernatant of LPS-stimulated (10 ng/mL concentration) PBMC after overnight incubation (**B**–**D**). Secreted IL-1β (B), TNF (**C**), and IL-6 (**D**) cytokines were quantified by flow cytometry and are shown as % of stimulation concentrations of the LPS condition (100%, red dotted line). Bar plots show each experimental group’s mean normalized cytokine expression (±SEM); each point represents the replicates (n = 3). *\*\*\*\*p* ≦ 0.0001, \*\**p* ≦ 0.0021, \**p* ≦ 0.0332, one-sample *t*-test.

Prior to anti-inflammatory screening using *ex vivo* human peripheral blood mononuclear cells (PBMC) and *in vivo* animal model (mice), we conducted *in vitro* toxicity testing for compounds **1–4** against bile duct epithelial (H69) cells using real-time cellular monitoring - an xCELLigence system (SI Appendix, Fig. S15). None of the four tested compounds was toxic to cells at a 10 μg/mL dose concentration (see Fig. 1A), so they were further evaluated for their anti-inflammatory activity using standardised *in vitro* and *in vivo* models of IBD.

### 3.1 Compounds 2–4 significantly suppressed the release of colitis-associated pro-inflammatory cytokines by human PMBCs

The anti-inflammatory activity of non-toxic compounds (**1**–**4**) was evaluated using human PBMCs, which contain various immune cells, including lymphocytes, monocytes, and natural killer (NK) cells(Luna et al., 2010). We used LPS to stimulate innate immune cells and showed that compounds **2**, **3**, and **4** (at 10 µg/mL concentration) significantly suppressed levels of IL-1β (Fig. 1B). Moreover, compounds **2** and **3** also significantly suppressed levels of TNF (Fig. 1C) and IL-6 (Fig. 1D). Tested compounds did not suppress the anti-inflammatory cytokine IL-10, which was good (SI Appendix, Fig. S15). Lipopolysaccharide explicitly activates monocytes and macrophages, but not T-cells(Meng and Lowell, 1997). Since compounds **2**–**4** significantly suppressed the release of colitis-associated pro-inflammatory cytokines by human PMBCs, we further investigated their anti-inflammatory potential in the trinitrobenzene sulfonic acid (TNBS)-induced acute colitis mouse model.

### 3.2 Compounds 3 and 4 protected mice against TNBS-induced colitis

Based on the *in vitro* cytotoxicity (Fig. 1A and SI Appendix, Fig. S15) and anti-inflammatory results (Fig. 1B-D), compounds **2**–**4** were further investigated for their ability to protect against TNBS-induced acute intestinal inflammation in mice. TNBS induces intestinal inflammation characterised by Th1 and Th2 cytokines, and causes gut inflammation that resembles clinical and histopathological symptoms experienced by patients with Crohn’s Disease and Ulcerative Colitis (UC)(Shepherd et al., 2018). Of the three compounds tested, mice treated with 20 μg of compound **3** (garcinia biflavonoid-1) and compound **4** (parvifoliol F) showed significant improvement in clinical symptoms or clinical score (such as body weight loss, piloerection, mobility, fecal consistency), macroscopic score based on colon shortening, adhesion, edema, and ulceration; and histopathology score including colon wall and lamina propria thickenings, crypt damage/goblet cells, necrosis and inflammatory cell infiltrations (Fig. 2A-E)) compared to the ‘TNBS’ sick control group (Fig. 2A). Considering these overall disease outcomes, compound **4** significantly alleviated colon inflammation. Compound **3** also showed an improvement in the disease outcomes but as statistically significant as compound **4**. The hematoxylin and eosin-stained (H&E) paraffin-embedded colon sections (Fig. 2E) were examined to assess histological changes induced by TNBS administration. The “TNBS” group displayed severe lesions and significant damage to the gastric mucosa (labelled ‘**b**’ in Fig. 2E–TNBS) compared to naïve/healthy control mice, which had healthy colons. The group treated with compound **4** (Fig. 2E) had colon architecture with intact lamina propria, epithelial compartments, and healthy goblet cells comparable to the Naïve group (labelled ‘**a**’ in Fig. 2E–Naïve). Moreover, compound **4** significantly reduced proinflammatory cytokines, IL-1α and IL-6, in mouse colon homogenates (Fig. 2F&G). Compound **3** also showed a similar trend of reduced IL-1α and IL-6 levels, but these values did not reach statistical significance (*p* > 0.05).

**Fig. 2.**
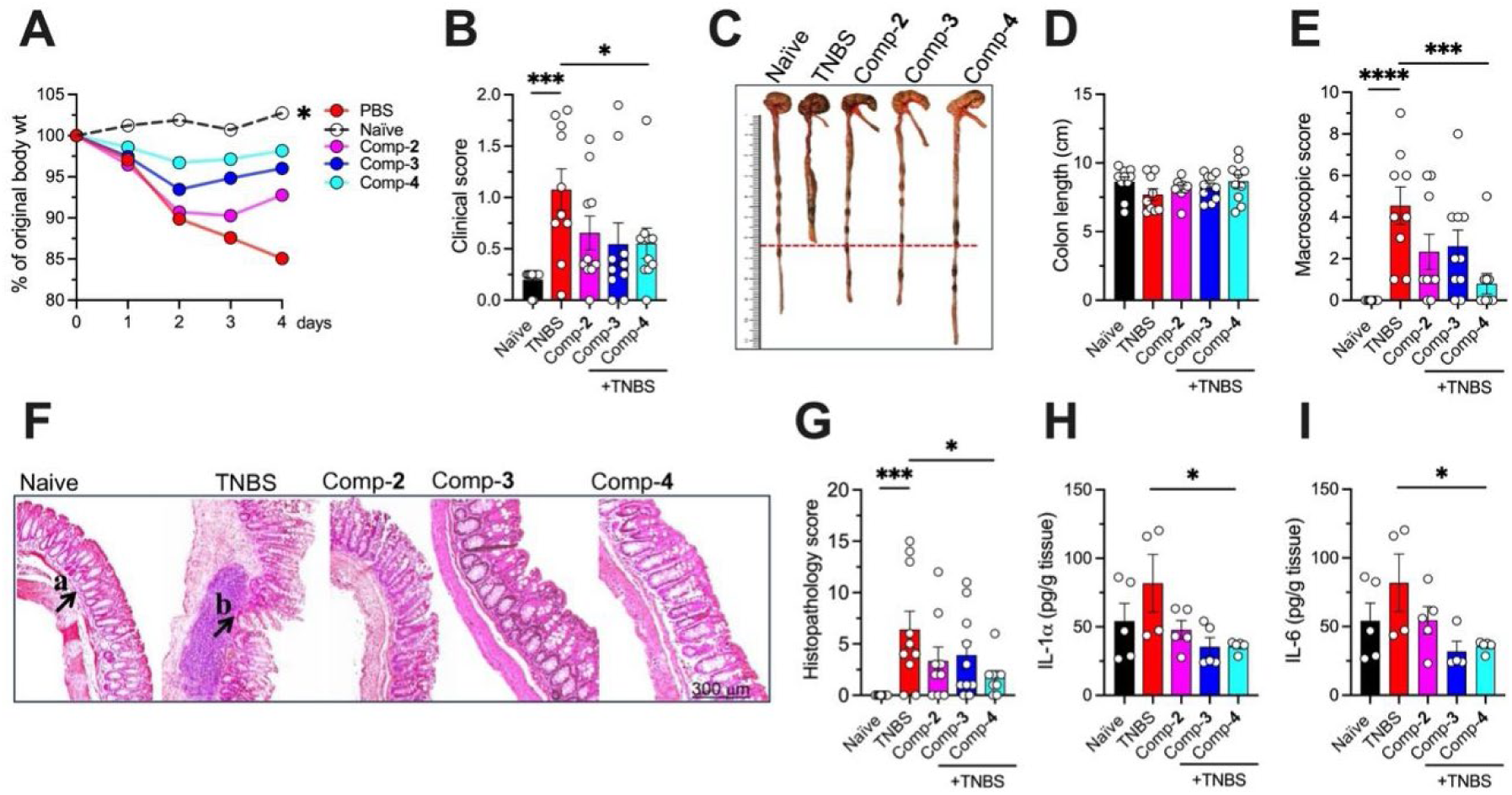
Testing of anti-inflammatory compounds using the *in vivo* TNBS mouse model of colitis (*n* = 10 per group). BALB/c mice were induced colitis on day zero by administering 2.5 mg/100 μl TNBS in 50% ethanol intrarectally. The next day, mice were treated with 20 μg of anti-inflammatory compounds (**2**, **3**, and **4**). Control mice received an equal volume of PBS vehicle. (**A**) Mean percent change of initial body weight, * shows the statistical significance observed on the fourth day of the experiments. (**B**) Clinical scores (mean and individual data points) consist of fecal consistency, motility, and piloerection scores. (**C**) Macroscopic colon pathology scores (mean and individual data points) at day-4 necroscopy. (**D**) Histopathological scores of hematoxylin and eosin-stained (H&E)-stained colon sections (Mean and individual data points). (**E**) Representative photos of H&E-stained colon sections. The arrow indicates the epithelial cell damage and the infiltration of immune cells. (**F** & **G**) Levels of proinflammatory cytokines IL-1α and IL-6 were significantly reduced in colon homogenates of treated mice (n = 5 mice per treatment group). \**p* ≦ 0.05, \*\*\**p* ≦ 0.001, \*\*\*\**p* ≦ 0.0001, Mann-Whitney *U* test compared to PBS control. H&E-stained images were captured using an APERIO CS2 scanner (Leica Biosystems Imaging, Inc., USA) at 40× magnification.

### 3.3 Compounds 3 and 4 promoted gut microbiome homeostasis in mice with TNBS-induced colitis

Several studies have hypothesized and shown a strong association between IBD and dysbiosis in gut microbiota. Thus, we investigated whether the improvement in colon morphology of treated mice could be due to the efficacy of compounds in protecting against gut dysbiosis caused by TNBS-induced colitis. 16S rRNA sequencing demonstrated that both compounds restored gut microbial homeostasis by reversing colitis-associated dysbiosis and reducing inflammation-associated microbial signatures. Mice treated with compound **3** showed significantly higher alpha diversity than the naive group (Fig. 3A), which had the lowest bacterial richness and diversity. Both Shannon and Simpson indices were elevated in the group treated with compound **3**, while the group that received compound **4** showed intermediate values, lower than the vehicle-treated group but higher than naïve mice that did not receive TNBS. Observed richness was highest in the group that received compound **3**. Beta diversity analysis (Fig. 3B) indicated that the gut microbiota of the naive group clustered more closely with compound **3** and **4** groups, while the TNBS group had a distinct cluster closer to the compound **3** group.

**Fig. 3.**
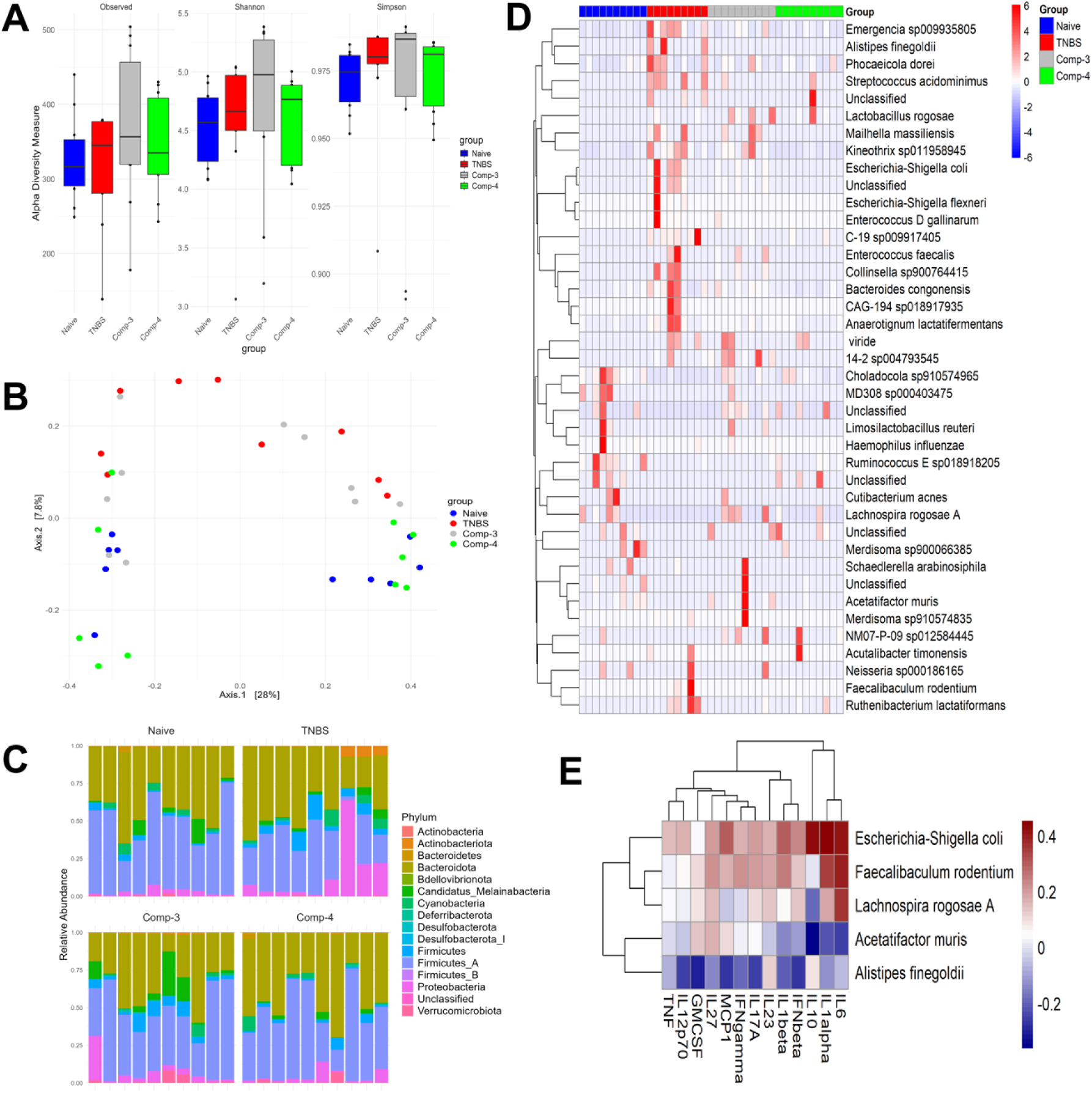
Gut microbial diversity and community composition in the TNBS-induced colitis mouse model. (**A**) Alpha diversity was evaluated using the observed richness, Shannon, and Simpson indices, capturing microbial diversity by assessing species abundance and distribution. Box plots demonstrate the variability across the naive, TNBS, compound **3**, and compound **4** groups, revealing notable differences in microbial diversity. (**B**) Principal Component Analysis (PCA) showing beta-diversity of microbiome community based on Bray-Curtis ordination metrics shows the distinct clustering of microbial communities within each group, providing insights into structural differences in the gut microbiota. (**C**) Relative abundance plots illustrate the phylum-level variations in microbial composition across the four groups. (**D**) Heatmap of species-level abundance highlights the top 20 bacterial species across groups (Padj<0.05), showcasing the species distribution and their respective abundances (for details, please see SI Appendix, Fig. S17 and Table S3). (**E**) Spearman’s correlation heatmap to explore the relationships between dominant microbiota at the species level and the cytokines measured in this study.

The phylum-level relative abundance analysis showed that *Firmicutes* and *Bacteroidetes* were the predominant phyla across all groups (Fig. 3C). However, the abundance of *Firmicutes* was significantly reduced in the TNBS group. Compared to the naive group, the TNBS group exhibited a marked increase in *Proteobacteria* populations alongside a significant decrease in *Firmicutes* (*p* < 0.01). Notably, mice treated with TNBS and compounds **3** and **4** showed a significant reduction in *Proteobacteria* and a corresponding increase in *Firmicutes* (*p* < 0.05), with minimal changes observed in the abundance of the *Bacteroidetes* phylum.

Previously, clinical studies have shown that UC patients exhibit a lower *Firmicutes* to *Bacteroides* ratio (F/B)(Stojanov et al., 2020), while mice with colitis demonstrate a significantly higher abundance of *Proteobacteria*, a hallmark of gut dysbiosis(Boger-May et al., 2022). In our study (Fig. 3D and SI Appendix, Fig. S17 and Table S3), we identified two species belonging to the *Firmicutes* phylum, *Lachnospira rogosae* A and *Acetatifactor muris*, which were abundant in naive mice but significantly decreased in the TNBS-induced colitis mice (*L. rogosae* A: -26.03 log2 fold change, *p* = 1.79E-18; *A. muris*: -26.35 log2 fold change, *p* = 2.16E-21). Remarkably, TNBS-induced colitis mice treated with compounds **3** and **4** led to a significant increase in *L. rogosae* A abundance (*L. rogosae* A: 26.52 log2 fold change, *p* = 2.46E-19 for compound **3**; 21.86 log2 fold change, *p* = 1.85E-13 for compound **4**). Similarly, *A. muris* abundance was significantly restored (26.67 log2 fold change, *p* = 1.07E-23) when the mice were treated with compound **3**. Additionally, we observed a significant increase in the abundance of *Escherichia-Shigella coli* species, part of the *Proteobacteria* phylum, in the TNBS-induced colitis group (16.90 log2 fold change, *p* = 7.78E-33). Interestingly, mice treated with compounds **3** or **4** had a sharp reduction or even complete elimination of pathogenic *E.-Shigella coli*.

Additionally, a few other bacterial species, such as *Facecalibaculum rodentium* (24.92 log2 fold change, *p* = 3.85E-17) and *Alistipes finegoldii* (24.96 log2 fold change, *p* = 4.35E-17), showed a significant increase in the TNBS-induced mice group. Interestingly, treatment with compounds **3** and **4** resulted in a marked reduction in the abundance of *F. rodentium* (-24.25 log2 fold change, *p* = 1.47E-17 for compound **3**; -24.76 log2 fold change, *p* = 6.10E-17 for compound **4**) and *A. finegoldii* (-24.69 log2 fold change, *p* = 7.55E-17 for compound **3**; -25.15 log2 fold change, *p* = 1.96E-17 for compound **4**). These findings suggest that compounds **3** and **4** effectively reduce the elevated abundance of bacterial species associated with TNBS-induced colitis, thereby promoting gut-microbiome symbiosis.

### 3.4 Gut microbial species are significantly associated with inflammatory cytokine responses

We conducted Spearman’s correlation analyses to explore the relationships between dominant microbiota at the species level and the cytokines measured in this study (Fig. 3E). All selected bacterial species showed high correlations with at least one of the cytokines tested. Notably, the abundance of *E.-Shigella coli* and *F. rodentium* exhibited a strong positive correlation with IL-1α and IL-6 responses (Spearman’s correlation coefficient = 0.4). Moreover, *E.-Shigella coli* abundance was also strongly linked to the production of several other pro-inflammatory cytokines, including IL-1β. In contrast, *Acetatifactor muris* abundance demonstrated a negative correlation with most of the cytokines, except IL-27 and IL-12p70. Similarly, *Alistipes finegoldii* abundance was negatively correlated with all observed cytokines except IL-23.

Functional impacts of gut microbiome alterations differed in mice treated with compounds 3 and 4. Although characterising compositional changes in the gut microbiota is essential, understanding the functional implications of these microbial alterations is equally important for interpreting their biological relevance. Therefore, functional pathway prediction was performed using PICRUSt2 based on full-length 16S rRNA gene sequencing data. As shown in Fig. 4 and SI Appendix, Fig. S18, a total of 163 MetaCyc metabolic pathways were identified across all experimental groups. Among these, several pathways were significantly modulated following treatment with the plant-derived compounds, and only pathways demonstrating statistically significant differences are discussed here.

**Fig. 4.**
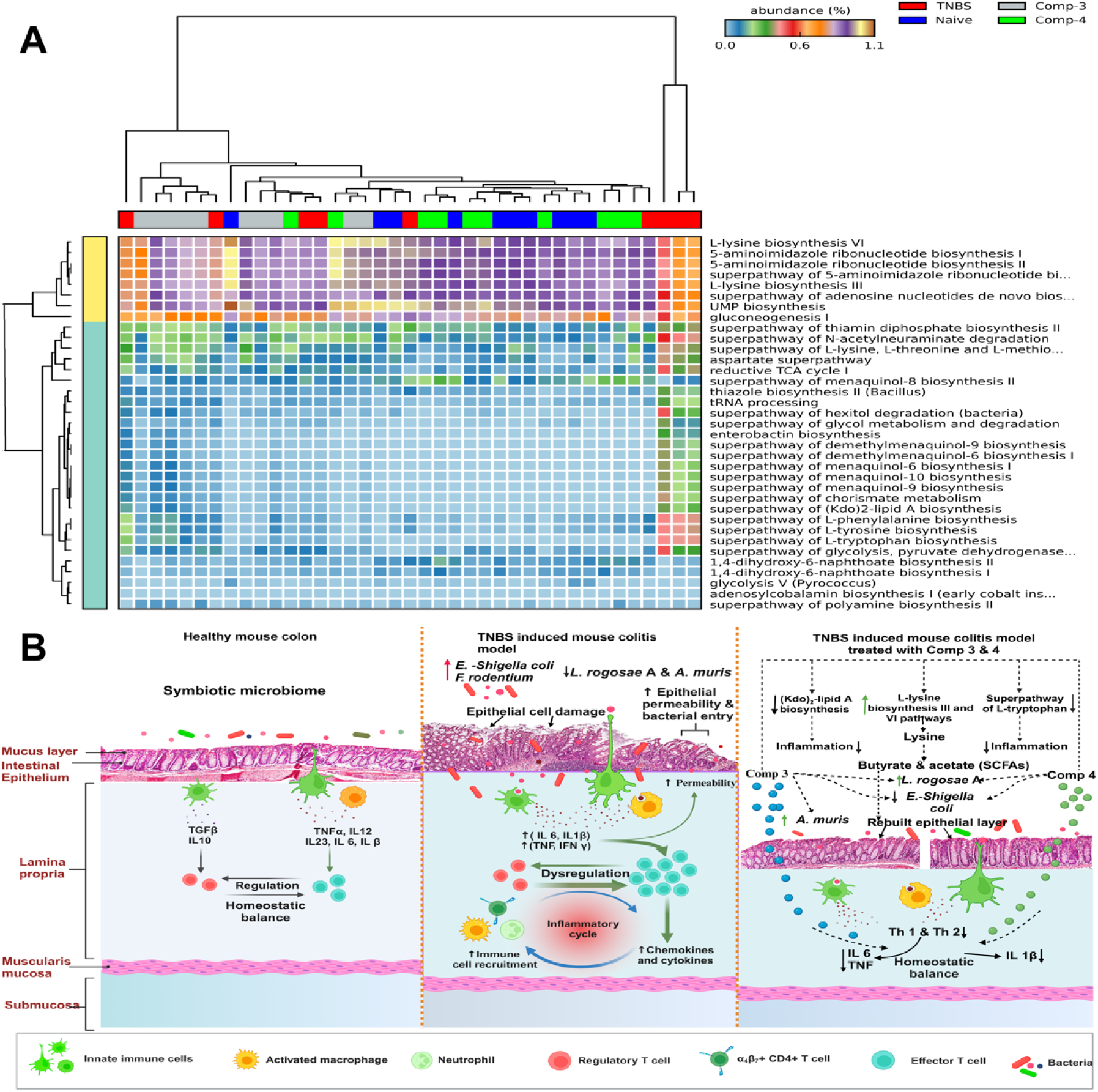
Functional prediction of the colon microbiota revealed using PICRUSt2. (**A**) Naïve vs model, (B) model vs. plant compound **3**, (**C**) model vs. plant compound **4**. Significantly altered microbial metabolic pathways predicted by PICRUSt2 are presented as extended error bar plots. Bar graphs on the left represent the mean relative abundance of each pathway, while dot plots on the right indicate differences in mean proportions between the compared groups based on Welch’s test (only pathways with *p* < 0.05 are shown). (**D**) Heatmap summarising top 35 pathways with *p* < 0.005 across different mouse groups (please see Figs. S18-S21 for group-wise enriched pathways). (**E**) Potential mechanism of action between plant compounds and the gut microbiota in a colitis mouse model.

Treatment with the plant-derived compounds substantially altered multiple microbial metabolic functions in TNBS-induced colitis mice. In particular, microbial pathways associated with L-lysine biosynthesis III and VI were significantly depleted in the TNBS group relative to healthy controls. Administration of compounds **3** and **4** significantly increased the abundance of these microbial populations. This observation is consistent with findings in the human gut bacterium *Intestinimonas* strain AF211, which can convert lysine into butyrate and acetate – key energy sources for enterocytes that help maintain colon health (Fig. 4A)(Bui et al., 2015). In addition, colitic mice displayed a notable increase in microbial populations related to the (Kdo)₂-lipid A biosynthesis and super pathway of L-tryptophan biosynthesis compared to the control group. Interestingly, compounds **3** and **4** significantly reduced these populations. The (Kdo)2-lipid A biosynthetic pathway is the hallmark of the extreme evolution of *Proteobacteria*, especially *Escherichia coli(Opiyo et al., 2010)*, which is closely associated with IBD. In addition, tryptophan metabolism in the gut is closely linked to immune regulation and inflammation, especially through activation of the kynurenine pathway. These results align with previous murine studies, where overactivation of the kynurenine pathway was strongly linked to heightened inflammatory responses(Michaudel et al., 2023). Furthermore, mice treated with compounds **3** and **4** showed marked decreases in microbial populations associated with several important metabolic pathways, including L-tyrosine biosynthesis, L-phenylalanine biosynthesis, and polyamine biosynthesis II, compared to colitic mice that received vehicle. Previous research has shown that inhibiting the polyamine biosynthesis pathway can reduce T-cell-mediated inflammation and alleviate inflammatory diseases(Wu et al., 2020).

### 3.5 LEGENDSCREEN™ identified differentially expressed IBD-related surface markers/genes in P/I-stimulated PBMC when treated with compound 3

Based on *in vitro* and *in vivo* data, compound **3** was the most promising. We further investigated compound **3** for its impact on T-cell responses by stimulating human PBMC with phorbol 12-myristate 13-acetate/ionomycin (P/I), as T cells play a vital role in sustaining IBD pathogenesis. Compound **3** significantly suppressed the secretion of TNF, IL-6, and IL-18 (Fig. 5A). To gain more insight into the drug-like properties of compound **3**, we further aimed to generate a robust human protein profile of P/I-stimulated PBMC enriched with leucocyte-specific transmembrane glycoprotein, CD45. CD45-enriched PBMC treated with compound **3** were stained for CD3+ and CD11c cells. Subsequently, these stained cells were subjected to a LEGENDScreen™ analysis using the LEGENDScreen (BioLegend) human cell screening (PE) kit. We used this kit because it includes 360 PE-conjugated monoclonal antibodies targeting cell surface markers linked to IBD. Therefore, it was expected to enable the identification of a wide spectrum of IBD-associated cellular markers following treatment. In total, 26 markers were differentially expressed in P/I-stimulated PBMC treated with compound **3** (Fig. 5B). Considering these 26 markers, the expression of protein markers was more for CD11c (antigen-presenting cells, APC – 21 markers) than CD3+ cells (T cells, 5 markers) (Fig. 5B). A total of 15 markers (12 on CD11c cells and 3 on CD3+ cells) were downregulated, and 11 markers were upregulated (9 on CD11c cells and 2 on CD3+ cells). For example, the chemokine receptor CXCR4 was downregulated on CD11c cells, and human leucocyte antigen (HLA), CXCR1, and CD198 were downregulated on CD3+ and CD11c cells treated with compound **3** (Fig. 5B). Moreover, compound **3** also upregulated CXCR1 and CXCR5 on CD3+ cells.

**Fig. 5.**
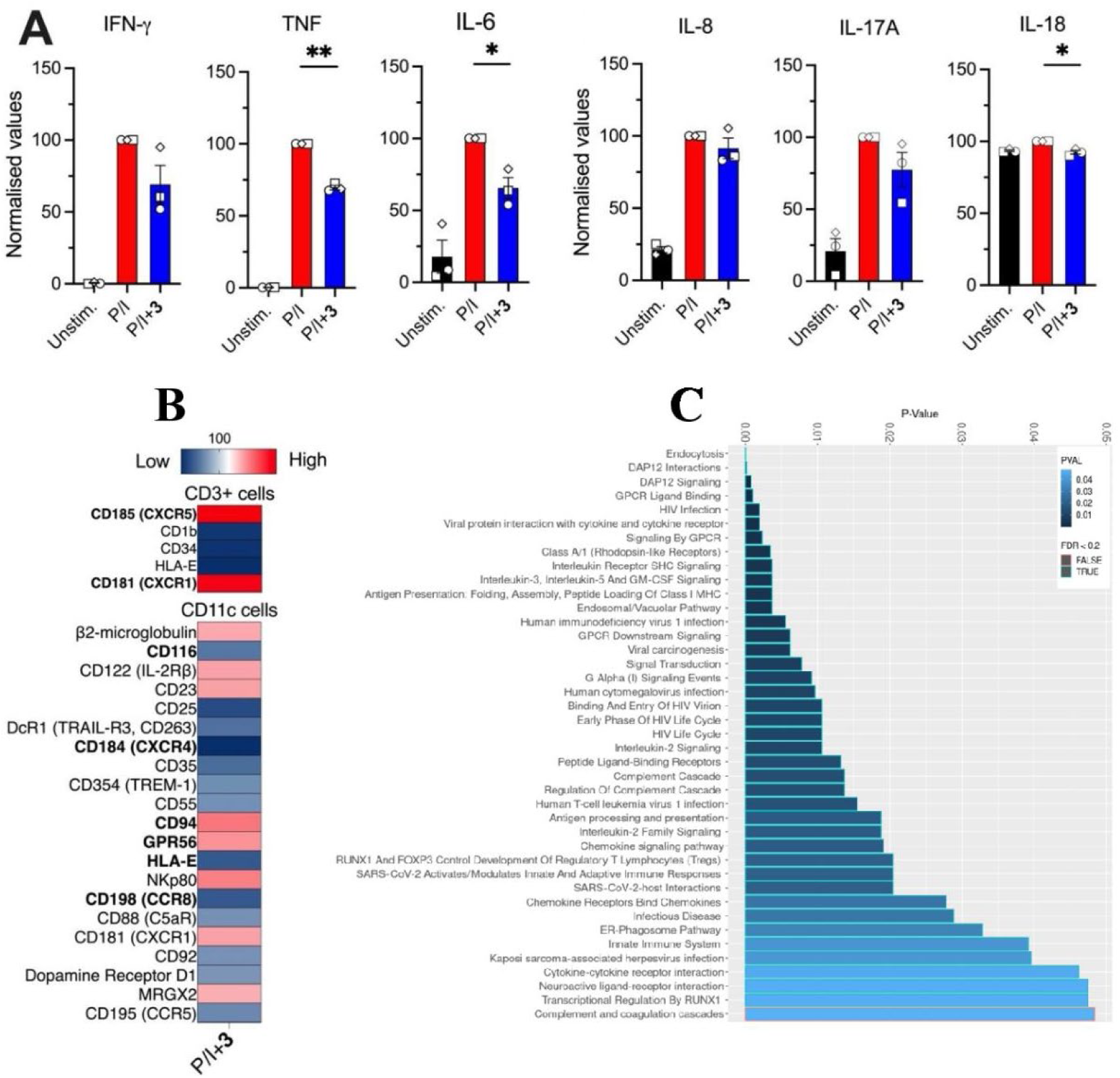
**(A) Anti-inflammatory effect of compound 3 (10 μg/mL concentration) was evaluated on PBMC stimulated with 1 μg/mL ionomycin and 50 ng/mL phorbol 12-myristate 13-acetate, P/I (n = 3 donors) after overnight incubation**. Secreted IFN-γ, TNF, IL-6, IL-8, IL-17A, and IL-18 cytokines were measured using flow cytometry and are presented as percentages relative to the stimulation levels observed under the P/I condition, which was set as 100% (red bar) for each donor. Bar plots show each experimental group’s mean normalized cytokine expression (±SEM); each point represents the donor mean of triplicate samples. \*\**p* ≦ 0.0021, \**p* ≦ 0.0332, one-sample *t-*test. (**B**) LEGENDScreen™ identified surface markers in human CD3+ and CD11c cell subsets. Human PBMC were isolated and enriched for CD45+ cells. Enriched cells were stained for CD3+ and CD11c+ cells. After collective staining, cells were submitted to LEGENDScreen™ and stained for 354 individual surface markers. Each marker’s mean fluorescence intensity (MFI) for each condition was normalized to the P/I-stimulated control to 100% and used as the baseline. Background gene lists for 354 markers were first input to DAVID and EnrichR using all xCELLigence panel targets found to correspond to annotated ENSEMBL genes. xCELLigence scores at least 1.5 times higher or lower relative to baseline levels were categorized as being up- or down-regulated, respectively. The heatmap represents up- or down-regulated markers, where PE expression ranged from low to high, with low (blue) and high (red). (**C**) Pathway enrichment analysis of markers differentially regulated by compound **3** in the CD11c cell subsets, as shown by LEGENDScreen™ analysis. For both DAVID and Enrichr pathway enrichment outputs, terms with a *p*-value of ≦ 0.05 were retained. A plot was generated with the R package ggplot for enriched pathways with sorted *p*-values on the y-axis and further annotated significantly enriched if the false discovery rate (FDR) *p*-value is < 0.2.

### 3.6 Pathway Enrichment Reveals Potential Anti-inflammatory Targets of Compound 3

Next, we conducted pathway enrichment analysis with markers differentially expressed by CD11c cells treated with compound **3** to shed some light on its possible anti-inflammatory mechanisms of action. CD3+ cell-expressed markers were excluded from the enrichment analysis as they were inadequate (only 5 markers were differentially expressed) for the analysis. In total, 40 pathways were significantly enriched (false discovery rate, FDR < 0.2) (Fig. 5C). The G protein-coupled receptors (GPCRs) signaling pathway, cytokine-cytokine receptor interaction, chemokine receptors binding chemokines, and chemokine signaling pathway are a few examples of pathways significantly enriched by compound **3** (Fig. 5C). Studies have shown the critical role of GPCRs in regulating inflammation and maintaining the gut membrane barrier and its significant association with the pathogenesis of IBD(Wasilewski et al., 2015; Zeng et al., 2020). Other pathways, such as Forkhead box P3 (FOXP3) protein and runt-related transcription factor 1 (Runx1) (Fig. 5C), that are important for the development of regulatory T cells (T_reg_ cells), particularly for their suppressive activity (Klunker et al., 2009), were also enriched in cells treated with compound **3.** Therefore, besides significant inhibitory activity induced by compounds **3** and **4** against proinflammatory cytokines, including IL-1β, IL-6, and TNF from *in vitro* and *in vivo* studies, our pathway enrichment analysis suggested that compound **3** may likely be interacting with G-protein coupled receptor (GPCR) (GPR35), FOXP3, and Runx1 transcription factors and the signaling pathways.

## 4. Discussion

This study characterized five compounds, including one novel natural product, a *p*-coumaric acid derivative called garcitine (4-*O*-β-*D*-Glucuronyl-*p*-coumaric acid) (**1**), along with four known compounds (**2**–**5**) from the aerial parts of *G. brassii*. These compounds, particularly **3** and **4**, demonstrated potent anti-inflammatory properties by reducing proinflammatory cytokines associated with IBD, including IL-6, both *in vitro* and *in vivo* settings. Interleukin (IL)-6 is pleiotropic, mediating host defense, immunity, disease pathogenesis, and inflammation(Narazaki and Kishimoto, 2018). Both IBD patients and chemically induced colitis mice showed increased production of IL-6 by CD4^+^ T cells and lamina propria macrophages(Atreya et al., 2000; Kai et al., 2005). In parallel, our LEGENDScreen profiling revealed upregulation of several T-cell surface markers, including the HLA class II molecule HLA-DRB1, a well-established genetic risk factor for IBD(Ashton et al., 2019; Venkateswaran et al., 2018). Notably, compound **3** also enhanced expression of chemokine receptors CXCR1 and CXCR5 on CD3⁺ T cells—receptors implicated in neutrophil recruitment and lymphoid tissue organization, respectively, both of which are dysregulated in IBD(Carlsen et al., 2002; Ishimoto et al., 2023; Legler et al., 1998; Williams and Parkos, 2007).

In addition to immunomodulatory effects, these compounds contributed to the restoration of gut microbial homeostasis, particularly by correcting dysbiosis within the *Firmicutes* and *Bacteroidetes* phyla. Integrative analysis of microbiome and cytokine profiles revealed associations between IL-1β/IL-6 levels and microbial taxa such as *E.–Shigella coli* and *F. rodentium*, underscoring a functional link between microbial composition and immune signaling in the inflamed gut. While previous studies have focused on host or environmental contributions to cytokine dysregulation(Li et al., 2016; Ter Horst et al., 2016), our findings highlight the potential of defined natural products to modulate both microbiota and host immune responses, offering a promising therapeutic avenue for IBD.

Despite increasing recognition of the link between gastrointestinal disorders and gut microbiome dysbiosis, it remains unclear which specific microbial members are responsible for triggering disease conditions, including IBD(Dixit et al., 2021). Dysbiosis in both the diversity and abundance of bacterial taxa, as well as in the gut metabolic profile, can serve as biomarkers for various diseases, including IBD(Carding et al., 2015). In this study, we investigated the pathological and immunological effects of plant-derived compounds, and compounds **3** and **4** demonstrated their potential to restore colon morphology (Fig. 2E) by modulating dysbiotic gut microbiota. In our TNBS-induced colitis mouse model, IBD-associated bacteria such as *E.-Shigella coli*, *Alistipes finegoldii*, and *F. rodentium* were enriched (Fig. 3D), while several beneficial bacteria from the dominant phylum *Firmicutes* were significantly depleted (Fig. 3C & D). Treatment with plant-derived compounds (**3** and **4**) effectively restored these dysbiotic bacterial species, promoting gut microbiome symbiosis and potentially alleviating colitis.

Understanding the interplay between gut microbiota and the immune system is crucial for unraveling immune-mediated and infectious diseases. For instance, variations in the abundance of certain gut bacteria have been linked to cytokine responses, potentially affecting disease susceptibility(Kau et al., 2011; Yoo et al., 2020). This influence appears to be exerted on the intrinsic capacity of immune cells to produce cytokines, rather than altering the number of circulating immune cells. Our analysis reveals a significant link between microbial abundance and cytokine response variability (Fig. 3E). Notably, the elevated abundance of *E.-Shigella coli* and *F. rodentium* was found to be strongly positively correlated with several proinflammatory cytokines, including IL-1α, IL-1β, and IL-6. Conversely, the abundance of *Lactobacillus rogosae A* and *Acetatifactor muris*, both from the phylum *Firmicutes*, displayed a negative correlation with most proinflammatory cytokines assessed in this study. This observation aligns with an expanding body of research suggesting that *Firmicutes* play a role in regulating systemic immunity(Jordan et al., 2023), promoting overall physiological balance. These findings underscore the immune system’s precise recognition and interaction with microbial species and their metabolites, highlighting the specific contributions of distinct microbial factors to immunological responses.

Our findings reveal that plant-derived compounds (**3** and **4**) significantly influenced several gut microbial metabolic pathways in TNBS-induced colitis mice (Fig. 4A). The increased activity in L-lysine biosynthesis III and VI pathways following treatment aligns with previous studies on amino acid metabolism, especially lysine, which plays a role in producing bioactive products by several *Bacteroides* spp. and some *Firmicutes(Ou et al., 2013; Yang et al., 2023)*. These products, particularly butyrate, are essential for maintaining colon health by serving as an energy source for colonocytes(Bui et al., 2015). Additionally, our observation of reduced (Kdo)₂-lipid A biosynthesis and the superpathway of L-tryptophan biosynthesis highlights the potential role of plant compounds in modulating gut microbial activities linked to IBD pathogenesis. *E. coli*, a significant contributor to IBD, evolves through the (Kdo)₂-lipid A pathway(Opiyo et al., 2010), while the kynurenine pathway of tryptophan metabolism is well-known for its association with inflammation(Rahaman et al.). Our results are consistent with murine models showing that overactivation of this pathway is linked to heightened inflammatory responses. Moreover, the observed decrease in microbial populations associated with polyamine biosynthesis II, L-tyrosine, and L-phenylalanine pathways suggests these compounds may attenuate inflammation by suppressing T-cell-mediated responses. Polyamine biosynthesis is crucial for immune regulation, and its inhibition has been shown to alleviate inflammatory conditions (Wu et al., 2020). Taken together, our results support the hypothesis that plant-derived compounds modulate gut microbial composition and metabolism in ways that reduce inflammation (Fig. 4B). These insights align with the growing body of evidence suggesting that targeting gut microbiota could provide therapeutic benefits for IBD patients.

## 5. Conclusion

In summary, this study provides compelling evidence that the plant-derived compounds **3** (garcinia biflavonoid 1) and **4** (parvifoliol F) alleviate inflammation via downregulation of proinflammatory cytokines and restoration of beneficial microbiota in the experimental colitis mouse model. The observed reduction in proinflammatory cytokines, including IL-6, and the restoration of symbiotic gut microbiota in the TNBS-induced colitis model (Fig. 3E) highlight the potential therapeutic role of these natural products in addressing gut dysbiosis and immune imbalance associated with IBD. Moreover, compound **3** can also assist in regenerating the microbiome without altering the natural gut environment in the experimental mice. Thus, by influencing key metabolic pathways and bacterial species linked to inflammation, compounds **3** and **4** not only demonstrate their efficacy in alleviating colitis but also underscore the potential of targeting the gut microbiome for future IBD treatments. Further research is warranted to elucidate the exact mechanisms by which these compounds mediate their effects and to explore their potential in clinical applications for IBD management.

## Data availability

All study data are included in this article or provided as supplementary information. All raw sequencing data have been uploaded to the NCBI Sequence Read Archive (SRA) under the accession number SRP564249 and BioProject accession number PRJNA1223484.

## Acknowledgements

We thank Kim Miles (AITHM/JCU) for isolating human peripheral blood mononuclear cells (PBMC) and the Australian Institute of Tropical Health and Medicine (AITHM) for providing the laboratory facilities. This research was supported by the National Health and Medical Research Council (NHMRC) Ideas Grant (APP1183323) to P.W. and James Cook University postgraduate research scholarship (JCUPRS) to K.Y. S.S. is the recipient of an Australian Research Council Discovery Early Career Researcher Award (grant number DE200100367) funded by the Australian Government. The Australian Government had no role in study design, data collection, analysis, the decision to publish, or manuscript preparation.

## Author contributions

P.W. and D.C. conceptualized and designed the research; D.C., P.W., and K.Y. collected plant materials, and D.C. identified the plants; K.Y. isolated compounds, analyzed data, and wrote the original manuscript; N.L.D. and D.W. collected NMR data; S.G.P. P.W. and K.Y. characterized structures of isolated compounds; R.R., P.G., M.J.S., and K.Y. performed immunoassays and animal experiments; S.S. performed microbiome study, interpretation and wrote the original draft; M.Z.I. and M.R. performed microbiome study and data analysis; M.F., conducted gene ontology (GO) term and pathway enrichment analyses; P.W. acquired funding and resources involved in this research; AL., D.C., D.W., N.L.D., P.G., P.W., R.R., and S.S. edited the manuscript.

